# Large serine integrases utilise scavenged phage proteins as directionality cofactors

**DOI:** 10.1101/2024.08.21.608926

**Authors:** Abdulrazak Alsaleh, Tania Pena Reyes, Aron Baksh, Oluwateniola T. Taiwo-Aiyerin, Alexandria Holland, Ying Pigli, Phoebe A. Rice, Femi J. Olorunniji

**Affiliations:** School of Pharmacy & Biomolecular Sciences, Faculty of Science, Liverpool John Moores University, Byrom Street, Liverpool L3 3AF, U.K; Department of Biochemistry and Molecular Biology, The University of Chicago, Chicago, IL 60637, USA

## Abstract

Recombination directionality factors (RDFs) for large serine integrases (LSIs) are cofactor proteins that control the directionality of recombination to favor excision over insertion. Although RDFs are predicted to bind their cognate LSIs in similar ways, there is no overall common structural theme across LSI RDFs, leading to the suggestion that some of them may be moonlighting proteins with other primary functions. To test this hypothesis, we searched for characterized proteins with structures similar to the predicted structures of known RDFs. Our search shows that the RDFs for two LSIs, TG1 integrase and Bxb1 integrase, show high similarities to a single stranded DNA binding (SSB) protein and an editing exonuclease, respectively. We present experimental data to show that TG1 RDF is a functional SSB protein. We used mutational analysis to validate the integrase-RDF interface predicted by AlphaFold2 multimer for TG1 integrase and its RDF, and establish that control of recombination directionality is mediated via protein-protein interaction at the junction of recombinase’s second DNA binding domain and the base of the coiled coil domain.

## INTRODUCTION

Site-specific recombination events are frequently used for integration, excision, and inversion of mobile genetic elements in phages and prokaryotes (1–3). Typically, the process involves cleavage of all four DNA strands in recombining duplexes, exchange of DNA ends and re-ligation to give recombinant DNA products (4). A characteristic feature of reactions involving site specific recombination is the formation of covalent protein DNA intermediate, presumably to ensure that cleaved DNA ends are not lost during the strand exchange events. The identity of the nucleophilic amino acid that forms this covalent intermediate is the basis for classification of site specific recombinases as serine or tyrosine recombinases: Tyrosine recombinases e.g., Flp, Cre, and phage lambda integrases, utlise a catalytic tryrosine, while serine recombinases, e.g. large serine integrases (LSIs), many transposon resolvases and many invertases, use serine (4). The covalent intermediates store the energy of the broken phosphodiester bonds. Therefore, the net change in chemical bond energy for these reactions is zero, and therefore it cannot be used to drive the reaction forward (4)(Grindely 2006). Site – specific recombinase systems have evolved an array of different, sometimes complex strategies to favor the forward or reverse reaction direction (5–9). For LSIs, the balance between reaction directions is tipped by a second protein, termed a recombination directionality factor (RDF) (2, 10–12).

LSIs are used by temperate phages (and some mobile genetic elements) to insert their genomes into that of their host bacteria as stably integrated prophages (Figure 1). Integration involves recombination of short attachment sites (40-50 bp) on the phage (*attP*) and the bacterial host (*attB*) resulting in new sites *attR* and *attL* flanking the inserted prophage (2, 13). LSIs are typically phage-encoded proteins, and they form a group of highly conserved proteins that work via a similar mechanism (2).

**Figure 1:**
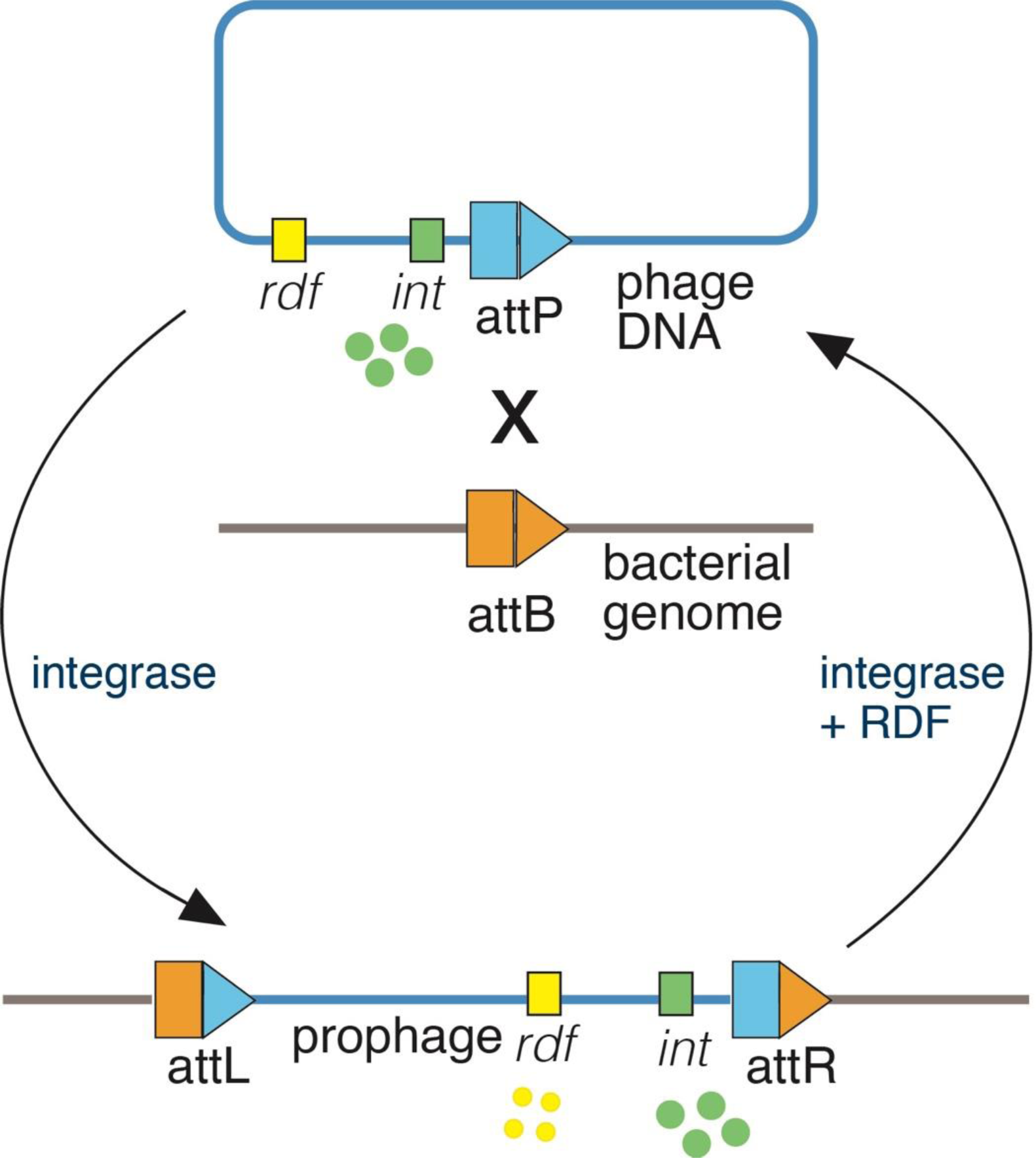
Phage integration and excision by large serine integrases. Phage-encoded LSI (green ovals) catalyse the integration of phage DNA at *attP* site (blue arrow) into the bacterial host genome at an *attB* site (amber). The recombination reaction results in the formation of recombinant *attR* and *attL* sites flanking the integrated prophage genome. In the reverse excision reaction, the recombination directionality factor, RDF (yellow ovals) is expressed from the prophage genome and binds to the LSI to promote *attR* x *attL* recombination.

Recombination of *attP* and *attB* is catalysed by the integrase in a synaptic tetramer complex in which dimers of integrase bind and synapse *attP* and *attB*, followed by DNA cleavage reactions, subunit rotation to exchange ends, and ligation of the recombinant DNA to form the products (*attR* and *attL*). In the lysogenic stage of the phage life cycle, the forward or ‘integrative’ reaction is irreversible resulting in stable insertion of the phage DNA into the host’s genome. The reverse ‘excision’ reaction happens during the lytic phase and requires an RDF to modify the preferred reaction direction of the integrase to promote *attR* x *attL* recombination. In addition to promoting excision, the RDF also inhibits integration to ensure the unidirectionality of LSI reactions either in the forward or reverse direction (10–12).

Tyrosine recombinases are also sometimes encoded by phages to function as integrases. However, they are more complicated: for example, phage lambda requires an ∼250bp attP site and additional host proteins (14–17). Furthermore, the mechanism for strand exchange by tyrosine integrases is different, as are their RDFs (often called “xis” proteins). RDFs for tyrosine recombinases have been shown to bind both DNA and the cognate integrase. However, some perform additional roles as transcriptional regulators (18).

Although LSIs can easily be identified in the genomes of phages (or mobile genetic elements), it is not readily obvious which genes code for RDFs. RDFs often lack synteny with their cognate LSIs, and there is no sequence homology across the set of known RDFs. Although there are no published experimental RDF structures, our recent analysis of predicted structures revealed no universally shared structural motifs that could explain their mechanism of action (19–22). However, they were all predicted to bind to the integrases at the same general location: the junction between the second DNA binding domain (DBD2; sometimes called the zinc-binding domain) and the coiled coil (CC) motif of the integrase. The conserved location of this interaction aided in developing a high throughput ‘virtual pulldown’ approach to finding RDFs for LSIs within phage genomes and suggests a conserved mode of action (19).

In contrast to the RDFs used by tyrosine recombinases, LSI RDFs generally have no known DNA binding role during recombination (11, 12, 18), and their functions appear to be solely to bind the integrase and modify its activity from integration-catalysing enzyme to one that promotes excision of the prophage from the host’s genome. The lack of DNA binding requirement may account for the diversity of proteins used by LSIs for RDF functions. This raises the question of how phages evolve functional RDF – LSI pairs. One likely mechanism is to repurpose existing proteins to acquire a moonlighting role as RDF. In fact, it has been shown that the phage Bxb1 RDF is indeed required for phage DNA replication as well as for excision (23). The tyrosine integrases encoded by phages HP1 and P2 use RDFs that also function as transcriptional repressors of early genes (24–27). Moonlighting is an approach used by phages since it allows them to derive broader functions using a limited genome (28–30).

We explored the possibility that RDF proteins for LSIs could have other ‘primary’ functions in the phage by searching for proteins with similar structures as the predicted structures for the known RDFs. We focused on the two largest known RDFs: that for phage Bxb1, and that for phage TG1 (which is also known as gp25, and which is similar to the RDFs for phages ϕC31 and ϕBT1 (31). Our findings show that these RDFs are different phage replication proteins moonlighting as RDFs for their cognate large serine integrases: the Bxb1 RDF is most likely the editing exonuclease subunit of its DNA polymerase, and the TG1 RDF is a bone fide single-stranded DNA binding protein.

## MATERIALS AND METHODS

### Structure predictions and bio-informatics

Predicted three-dimensional (3D) protein structures were initially predicted using the colabfold implementation AlphaFold2-multimer; version 1.5.2 with default parameters (20, 21, 32) (Figure 5). In the structures shown in Figure 5 the integrase sequence was truncated to include only the C-terminal segment, which includes DBD2 and the coiled coil. The inter- and intra-molecular measures of confidence for the AlphaFold2-multimer model of the TG1 RDF – integrase complex were ipTM = 0.71 and pTM = 0.5, respectively, and for the Bxb1 model were ipTM. = 0.80 and pTM = 0.65. RDF structures were later predicted with AlphaFold3 (22) in order to test addition of ions and ssDNA to the models (**Figure 2)**. The inter- and intra-molecular confidence scores for the Bxb1 model (RDF plus 1 Zn^2+^ ion, 1 Fe^3+^ ion and poly d(T)_6_) were ipTM = 0.86, pTM = 0.91, respectively, and for the TG1 model (RDF plus 1 Zn^2+^ ion and poly d(T)_6_) were ipTM = 0.82, pTM = 0.84. The structures were viewed and manipulated and figures made using PyMol (https://pymol.org/pymol.html). Surface electrostatic potential was calculated and visualized using the APBS PyMOL plugin (33).

**Figure 2:**
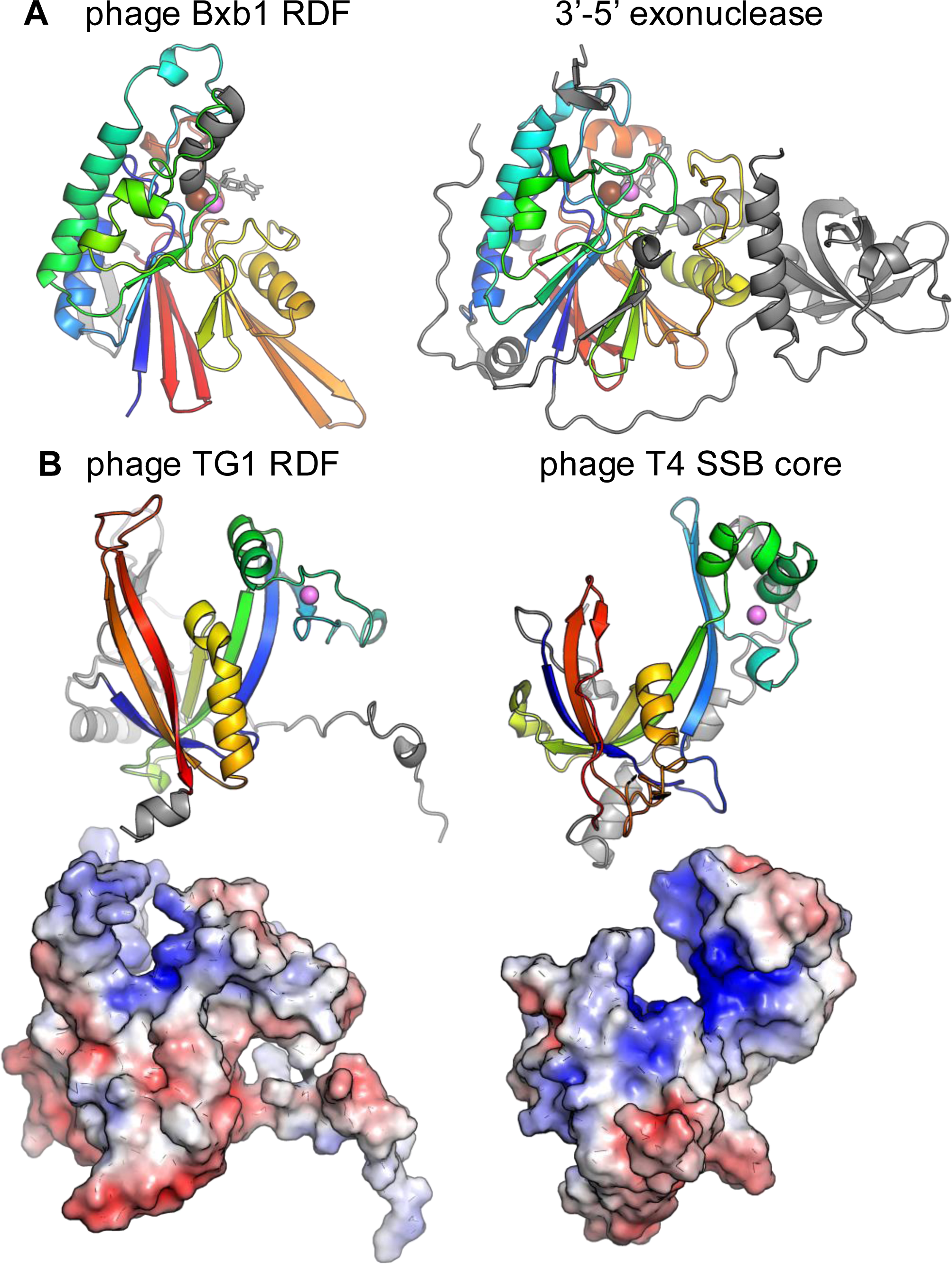
Structural similarities between Bxb1 and TG1 RDFs and replication proteins. (**A**) Comparison of the predicted Bxb1 RDF structure (left) to the crystal structure of the proofreading exonuclease subunit of PolD (right; PDB: 5IHE; (35). The related portions of each structure are shaded from N (blue) to C (red). Zinc ions are violet and iron ions brown. One nucleotide from the predicted structure of Bxb1 RDF with dT_6_ is shown in gray, as is the single nucleotide seen in the active site of the exonuclease. (**B**) Top: Comparison of the predicted TG1 RDF structure (left) to that of phage T4 single-stranded DNA binding protein core (right; PDB: 1GPC; (37). Bottom: Surface representations of each colored by electrostatic potential (blue positive, red negative).

The Dali server was used to find characterized proteins with structures similar to our predicted RDF structures (34). For the Bxb1 RDF, the top hits were all the small subunit of the D-family polymerase of Pyrococcus abyssi, which is an editing exonuclease with a purple acid phosphatase fold (35)– z scores for this subunit in different complexes ranged from 15.1 to 17.2. For the TG1 RDF, the search retrieved numerous OB fold domains, which are often used to bind ssDNA (36). The top hit was an ssDNA binding domain of BRAC2, with a z score of 4.0. However, it lacked the zinc-binding site predicted in the TG1 RDF. For this reason, for comparisons we choose to use the 5^th^ hit on the list (z score 3.3), gp32 from phage T4, a known single-stranded DNA binding protein that does carry a zinc-binding site (37). Although Alphafold3 predicted that ssDNA would bind in the positively charged cleft of both the TG1 RDF and phage T4 gp32, that is not shown in Figure 2 because there are no experimental structures showing ssDNA bound to T4 gp32.

### Integrase and RDF expression vectors for protein purification

Codon-optimised protein-coding DNA fragments expressing TG1 integrase and TG1 RDF were inserted into pET-28a(+) (Novagen), between the NdeI and XhoI sites as described previously (31, 38). Mutants of the integrase and RDF were made by cloning synthetic g-block DNA (Integrated DNA Technologies) containing the desired changes into the appropriate pET-28a(+)-based expression vectors. All proteins expressed from these plasmids carry N-terminal hexahistidine tags to allow purification via nickel affinity chromatography.

### *In vitro* binding reactions

The TG1 RDF protein used for *In vitro* binding reactions was purified as follows. E. coli (Rosetta(DE3)plysS) containing a pET vector encoding the TG1 RDF (described below) were grown at 37C in LB supplemented with (50 µg/ml kanamycin and 100 μM ZnSO_4_), induced by the addition of 0.5 mM IPTG once OD_600_ reach ∼1, then grown at 20 ^◦^C overnight (∼16 hours). Cell pellets were collected and resuspended in Ni-Buffer A (50 mM Phosphate, 1 M NaCl, 5% Glycerol, 1 mM TCEP, pH7.5) supplemented with complete Mini protease inhibitor cocktail (MilliporeSigma; 1 tablet per liter of culture). Lysozyme (200 ug/ml) was added before sonication. The sonicated sample was centrifuged at 20, 000 rpm in an SS-34 rotor for one hour at 4 °C, and the supernatant was collected and filtered. Affinity chromatography was carried out using a HisTrap HP column (Cytiva). The column was washed with Ni-Buffer A, after which the sample was loaded, washed with Ni-Buffer A, and eluted with Ni-Buffer B (Ni-Buffer A + 0.5 M imidazole, pH 7.5) following a 0–100% linear gradient over 30 minutes at a rate of 2 ml/min. Selected fractions were pooled and rechromatographed on the same column. Selected fractions were pooled again, then polished on HiPrep QFF ion exchange column (Cytiva) equilibrated with Q FF-Buffer A (20 mM Tris-HCl, 5% Glycerol, 1 mM TCEP, pH8). The column was then washed with 10% Q FF-Buffer B (Q FF-Buffer A + 2 M NaCl) after which the protein was eluted with Q FF -Buffer B (10-70% gradient; flow rate 2 ml/min). Fractions were chosen after SDS-PAGE and nuclease activity assay. To assay for nuclease activity, fractions were incubated with supercoiled plasmid (pUC19) and additional Mg^2+^followed by agarose gel electrophoresis. Purified protein was concentrated to 340 μM in 20 mM Tris pH 8, 0.5 mM EDTA, 200 mM NaCl, 20% Glycerol, 2 mM TCEP, flash-frozen in small aliquots in liquid nitrogen and stored at -80 ^◦^C.

DNA substrates for binding assays were purchased from IDT. The single-stranded substrate was poly(dT)_50_ with a 5’ fluorescein label (5’-/56-FAM). The 50 bp double-stranded substrate had a 5’ fluorescein label on the top strand, and the sequence: 5’ CAGCTCCGCGGGCAAGACCTAGCTCTTACCCAGTTGGGCGGGATAATTAA 3’. The binding assays were conducted in a total volume of 20 ul in buffer containing: 20mM Tris, pH 8, 100mM NaCl, 10% Glycerol, 50ng/μl BSA (bovine serum albumin), 1 mM TCEP. DNA substrates were present at 0.2 μM and protein at 0, 0.2 μM, 0.4 μM and 0.8 μM. After 1 hour incubation at room temperature, 15 μl of each sample was loaded onto a 10% 0.5 x TBE gel and run at 140 V for 2 hours at 4 ^◦^C. Bands were visualized on a Chemidoc imager. Due to the predicted zinc binding site in this protein, we also tested addition of 0.5 mM ZnSO_4_ but found that it did not change the DNA-binding ability of the protein.

### *In vitro* plasmid substrates for intramolecular recombination

The intramolecular recombination assay used is as described in Abioye *et al*. (38) and Olorunniji *et al.* (39). The plasmids for *attP* x *attB* and *attR* x *attL* recombination reactions are named pTG1-PBX and pTG1-RLX, respectively. In both substrates, the *att* sites are arranged in direct repeat or ‘head to tail’ orientation such that recombination results in resolution (excision) of the substrate plasmid into two separate smaller plasmid circles (Figure 4A). The sequences of the *attP, attB, attR, and attL* sites are shown in Figure 4B, and all the recombination substrate plasmids are available upon request. Supercoiled plasmid DNA used for *in vitro* reactions was prepared from transformed *E. coli* DS941 cells using a Qiagen miniprep kit (39). The concentrations of DNA preps were determined by measuring absorbance at 260 nm, and the quality of each prep was verified by electrophoresis on 1.2% agarose gels.

### Expression and purification of integrase and RDF for activity assays

Expression and purification of TG1 integrase and TG1 RDF, and their mutants were carried out as described in Olorunniji *et al.* (39) and Abioye *et al* (38). *E. coli* strain BL21(DE3)pLysS was made chemically competent and transformed with the appropriate protein expression vector. The expression strain for each protein was grown in 2x Y-broth at 37^◦^C to mid-log phase (OD_600_, 0.6 to 0.8) and cooled down rapidly to 20 ^◦^C before inducing protein expression with the addition of 0.5 mM IPTG, after which the cultures were grown for a further 16 hours at 20 ^◦^C. The cultures were grown with kanamycin (50 µg/ml) and chloramphenicol (50 µg/ml) added to the media. Cells were harvested by centrifugation at 4 ^◦^C, and the pellet was washed in 25 mM Tris–HCl (pH 7.5), 10 mM MgCl_2_, and the pellet was collected by centrifugation at 4 °C. The washed pellet was resuspended in 25 ml of Buffer A (20 mM sodium phosphate (pH 7.4), 1 M NaCl, 1 mM dithiothreitol (DTT) and 50 mM imidazole, 1% (v/v) ethanol). The suspension was cooled in ice, and the cells were lysed by sonication (Branson, SFX 150). The suspension was centrifuged for 30 minutes at 4 ^◦^C, 12 000 rpm, after which the supernatant was collected and filtered. Proteins were purified by nickel affinity chromatography using a HisTrap FF pre-packed column (GE Healthcare). The column was equilibrated with the starting Buffer A, at a constant flow rate of 1 ml/min, prior to loading the protein sample, also in Buffer A. The column was washed with Buffer A (25 ml) to remove unbound proteins and the bound protein of interest was eluted with Buffer B (Buffer A, but with 500 mM imidazole), increasing in a 0–100% linear gradient over 25 min. Purity of selected fractions was assessed by SDS-polyacrylamide gel electrophoresis and chosen fractions containing the protein of interest were dialysed against Protein Dilution Buffer (PDB; 25 mM Tris–HCl (pH 7.5), 1 mM DTT, 1 M NaCl and 50% v/v glycerol), and stored at -20^◦^C.

### *In vitro* recombination of supercoiled plasmid substrates, and product analysis

Purified integrases and RDFs were stored at -20 ^◦^C and diluted to the appropriate concentrations at 0 ^◦^C just before use. Integrases and RDFs are diluted in Protein Dilution Buffer as described above. *In vitro* recombination reactions were carried out as described in Abioye *et al*. (38). In summary, reactions were initiated by adding integrase (8 µM, 5 µl) to a 30 µl solution containing the appropriate substrate plasmid DNA (25 µg/ml), 50 mM Tris-HCl (pH 7.5), 100 µg/ml BSA, 5 mM spermidine, and 0.1 mM EDTA. For reactions involving integrase and RDF, equal volumes of integrase (16 μM) and RDF (16 μM) were mixed thoroughly and kept on ice for 15 minutes to ensure optimal protein-protein binding, then 5 μl of this mixture was added to the reactions. Reaction samples were incubated at 30 ^◦^C for 2 or 16 hours, after which the reactions were stopped by heating at 80 ^◦^C for 10 minutes to denature the proteins. The samples were cooled and treated with NruI (New England Biolabs) to facilitate separation and analysis of recombination products. This was done by mixing a 30 µl aliquot of the reaction mixture with 28 µl of B103 buffer (90 mM Tris-HCl pH 7.5, 20 mM MgCl_2_) prior to addition of 20 units (2 µl) NruI (New England Biolabs). The restriction digests were carried out at 37 ^◦^C for 2 hours. Following the digest, samples were treated with SDS and protease K by adding 7.5 µl loading buffer (25 mM Tris-HCl pH 8.2, 20%(w/v) Ficoll, 0.5% sodium dodecyl sulphate, 1 mg/ml protease K, and 0.25 mg/ml bromophenol blue) to the reaction sample and incubated at 50 ^◦^C for 30 minutes. The reaction products were separated by by electrophoresis on 1.2% agarose gels in 1x TAE, then stained with SYBR safe and visualised as previously described, using a BioRad GelDoc apparatus (38, 40). Digital images of the gels are shown in reverse contrast.

## RESULTS

### TG1 and Bxb1 RDFs are structurally homologous to DNA replication proteins

We recently showed that AlphaFold-predicted structures of serine integrase RDFs reveal a wide diversity of structures with no obvious pattern to explain their role as RDFs for the otherwise highly conserved LSIs (19). Despite this structural variation, most of the RDFs are relatively small proteins. The RDFs for SPbeta, A118, ϕRV1, Nm60, Bt24, Int10, and Int30 range between ∼7 and 9.3 kDa (19, 41–43). In contrast, RDFs of the ϕC31 family (ϕC31, ϕBT1, and TG1, ∼27 kDa) and Bxb1 (28 kDa) are larger proteins (11, 12); Figure 2 in (19)). However, for all of these RDFs, models of their complexes with their respective integrases predict interactions at the same integrase CC/DBD2 junction (19, 31). We hypothesized that the larger proteins in particular were likely to have been recruited to moonlight as RDFs and may have other biological functions.

Our search for characterized proteins with structures similar to those predicted for the two known types of large RDF – those for phages Bxb1 and TG1 - strongly supported the moonlighting hypothesis. Bxb1’s RDF was previously noted to have sequence homology with purple acid phosphatase-family enzymes, and to be required for phage DNA replication (10, 23). Its predicted structure is in agreement with that observation (Figure 2A), and our Dali search found that it is particularly closely related to the editing exonuclease subunit of some DNA polymerases (35). In contrast, the predicted fold of the RDF used by phage TG1 is similar to that of phage T4’s single-stranded DNA binding protein (SSB) (37) (Figure 2B). Furthermore, within each of these phages, the gene encoding the RDF is closer to other replication-related genes than to the integrase gene.

### TG1 RDF is a phage single-stranded DNA binding (SSB) protein

Based on the similarity between the predicted structure of TG1 RDF and the experimental structure of phage T4 SSB, we tested the ability of TG1 RDF to bind single-stranded DNA using an *in vitro* EMSA binding assay. The results show that the RDF binds single-stranded DNA in a concentration dependent manner (Figure 3). In contrast, the RDF does not bind double-stranded DNA.

**Figure 3:**
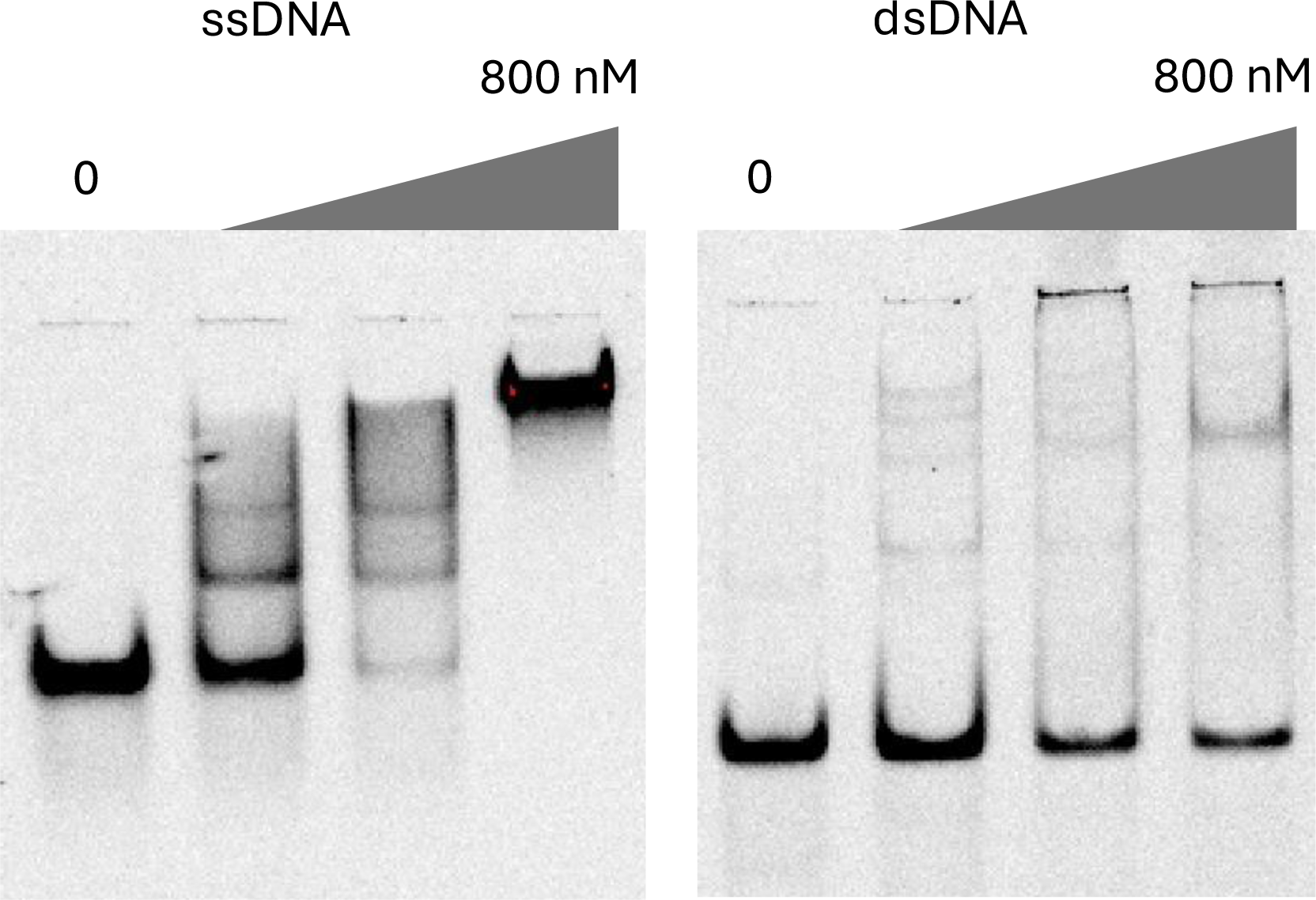
TG1 RDF is a single strand DNA-binding protein *in vitro*. Electrophoretic mobility shift assays of TG1 RDF (gp25) binding to ssDNA (left) and dsDNA (right). The protein was present at 0, 200, 400, and 800 nM and the 50 nt/bp fluorescein-labeled DNA substrate was kept constant at 200 nM.

Next, we asked if binding ssDNA could interfere with the RDF function of gp25 (Figure 4). As shown in Figure 4C, the presence of single-stranded DNA reduces the extent of activation of *attR* x *attL* reaction by the RDF. Correspondingly, single-stranded DNA relieves the inhibition of *attP* x *attB* reaction by the RDF (Figure 4D). In contrast, single-stranded DNA has no effect on *attP* x *attB* recombination by the integrase when the RDF is not included in the reaction (Figure 4E), showing that the effect of single-stranded DNA on the reactions seen in Figures 4C and 4D are due to interference with RDF function.

**Figure 4:**
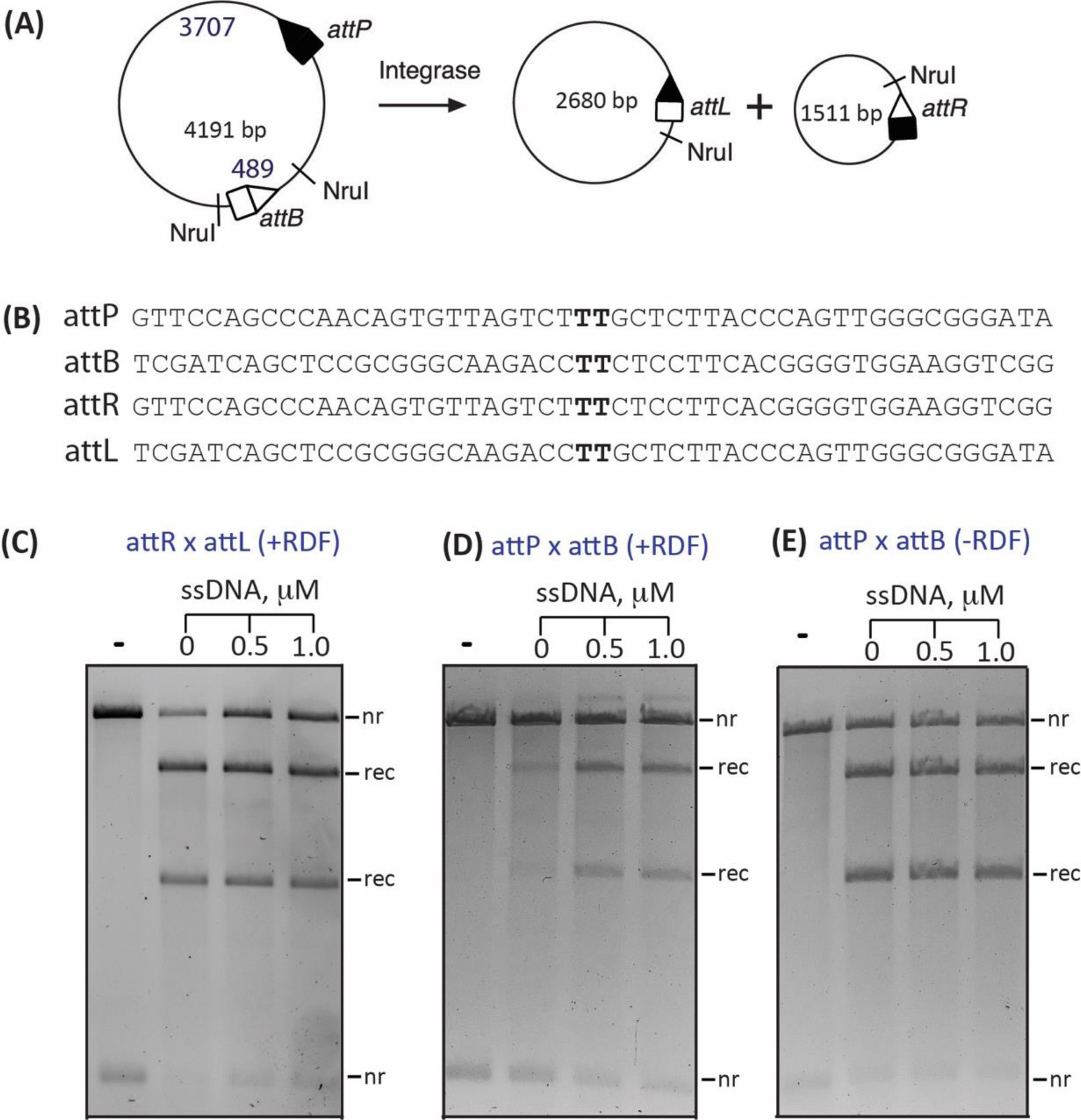
Single-strand DNA interferes with recombination functions of TG1 RDF in vitro. (A) Scheme depicting the in vitro intramolecular recombination reaction. Recombination of the plasmid substrate (pTG1PBX) gives two circular products in which the *attR* and *attL* sites are separated. For the *attR* x *attL* recombination reaction, the starting substrate plasmid has *attP* and *attB* sites replaced by *attR* and *attL* sites, respectively (pTG1RLX), with recombination giving *attP* and *attB* sites on separate circular plasmid products. (**B**) Sequences of recombination *att* sites for TG1 integrase. The effects of ssDNA on RDF function in in vitro recombination reactions are shown in C-E. (**C**) *attR* x *attL* in the presence of RDF and ssDNA, (**D**) *attP* x *attP* in the presence of RDF and ssDNA, and (**E**) *attP* x *attB* in the presence of ssDNA, without RDF. In all cases, reactions were carried out for 2 hours in the reaction buffer described in the Methods section (integrase, 800 nM; RDF, 800 nM), with the addition of 0, 0.5, and 1.0 µM ssDNA (polydT) as indicated. Reaction products were digested with the restriction endonuclease NruI prior to 1.2% agarose gel electrophoresis. The bands on the gel are labeled *nr* (non-recombinant, i.e. substrate), *rec* (recombination product).

### Experimental data support the modeled TG1 RDF - integrase interface

Next, we tested the functional relevance of the Integrase-RDF interface predicted by AlphaFold2-Multimer. We made specific amino acid changes on both the integrase and RDF (Figure 5) and tested the effects of the mutations on recombination activities *in vitro* (Figures 6 and 7). These mutations were spread across the predicted interface, and were designed to alter the charge in hydrophilic patches or to remove large hydrophobic side chains that could be important in the protein-protein interactions. As described below, the results of our mutational analysis are consistent with the model shown in Figure 5, and suggest that the portion of the interface highlighted in Figure 5E (involving the base of the integrase’s coiled coil (pale green)) is more important than the portion highlighted in Figure 5D.

**Figure 5:**
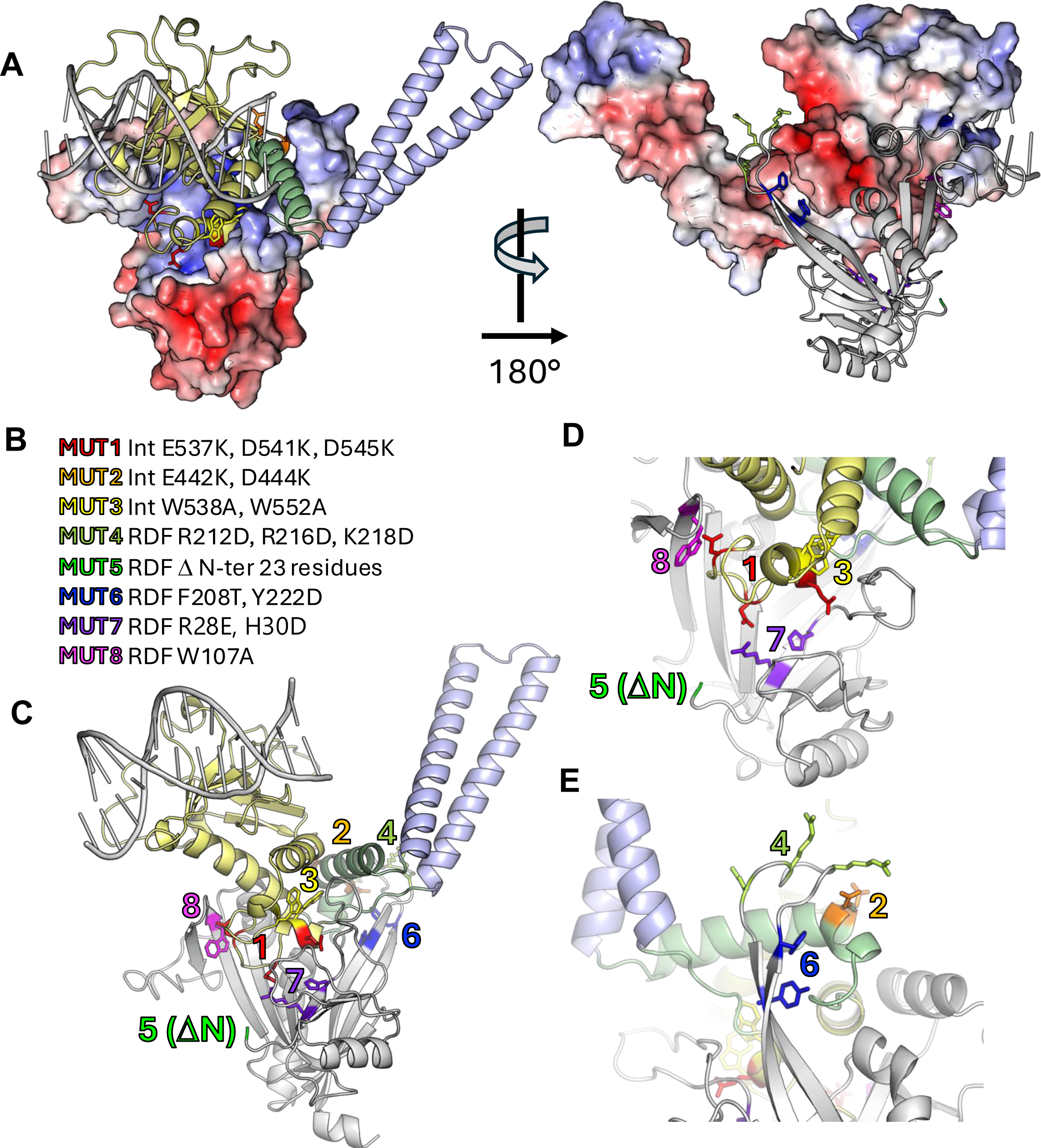
Integrase-RDF interface that mediates regulation of recombination. (**A**) Surface complementary of the predicted TG1 integrase -RDF interface. Left: The C-terminal domain of the integrase is shown as ribbons (DBD2, yellow; DBD2-proximal portion of the CC, pale green; CC pale blue), with DNA docked based on 4kis.pdb to guide the eye (48). A surface representation of the RDF is shown, colored by electrostatic potential (blue, positive; red, negative). Right: The RDF is shown as a gray ribbon, with the integrase surface colored by electrostatic potential. Side chains that were mutated are shown as colored sticks (see B). The C-terminus of the integrase (600-619) and the N-terminus (1–22) of the RDF are not shown as they were predicted to be disordered. (**B**), Table of mutations made. (**C),** The model shown in (A), with side chains mutated shown as sticks, color - coded as in B. **(D, E)**: close-up views of integrase – RDF interactions and the side chains that were mutated.

**Figure 6:**
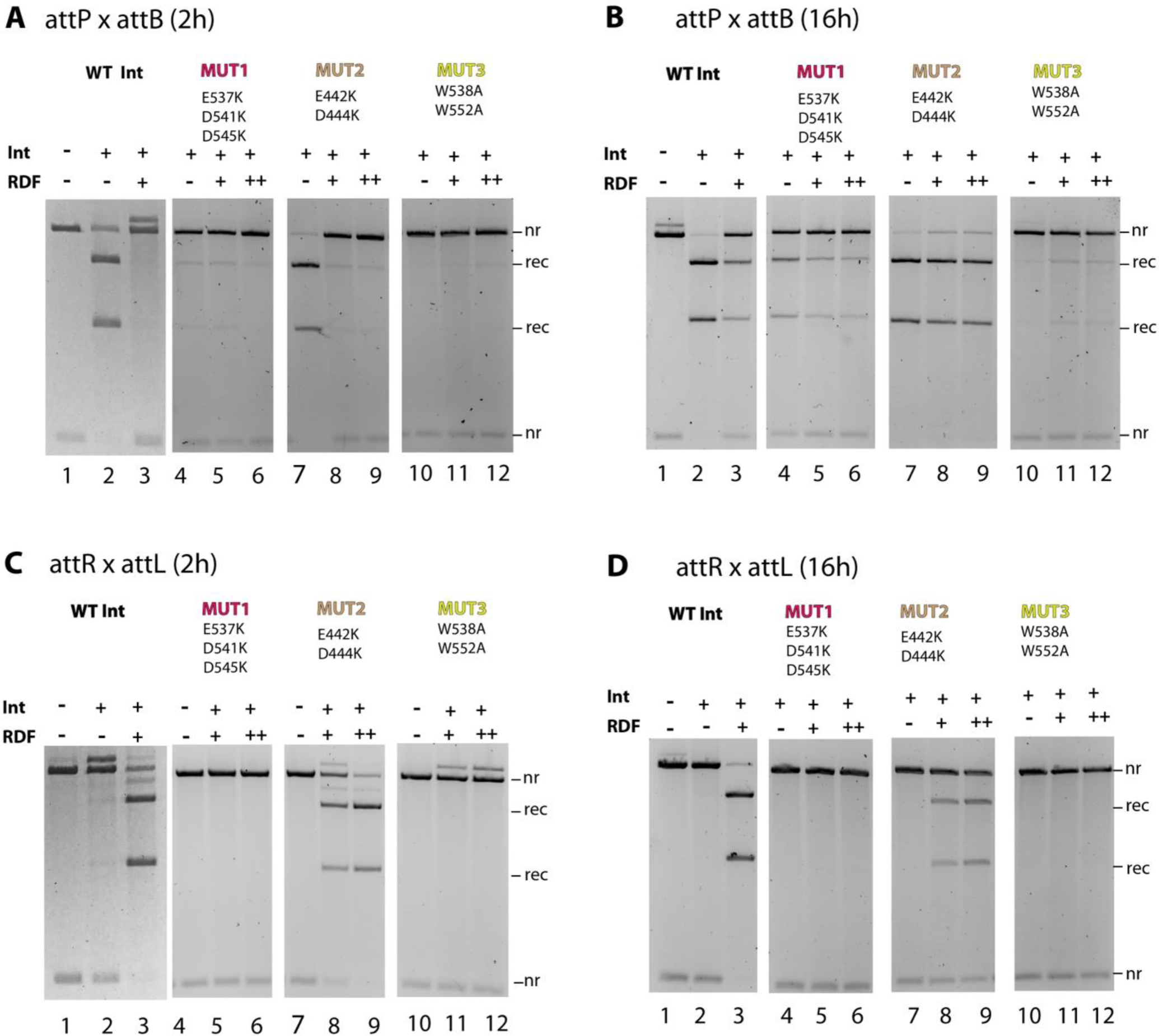
*In vitro* recombination activities of TG1 integrase mutants. Reactions were carried out for 2 or 16 hours in the reaction buffer described in the Methods section. **(A)** 2-h reaction, *attP* x *attB*. Integrase was not added to reaction in lane 1. **(B)** 16-h reaction, *attP* x *attB*. Integrase was not added to reaction in lane 1. **(C)** 2-h reaction, *attR* x *attL*. Integrase was not added to reaction in lane 1. **(D)** 16-h reaction, *attR* x *attL*. Integrase was not added to reaction in lanes 1. Reaction products were digested with the restriction endonuclease NruI prior to 1.2% agarose gel electrophoresis. The bands on the gel are labeled *nr* (non-recombinant, i.e. substrate), *rec* (recombination product). The final integrase concentrations in all reactions were 800 nM. The final RDF concentrations were 800nM (+) or 1600nM (++). The bands on the gel are labelled *nr* (non-recombinant, i.e., substrate) or *rec* (recombination product). The sizes of the products of the recombination reactions are shown alongside each band on the gel image. MUT1: TG1 Int-3K (E537K, D541K, D545K). MUT2: TG1 Int-2K (E442K, D444K). MUT3: TG1 Int-2A (W538A, W552A).

**Figure 7:**
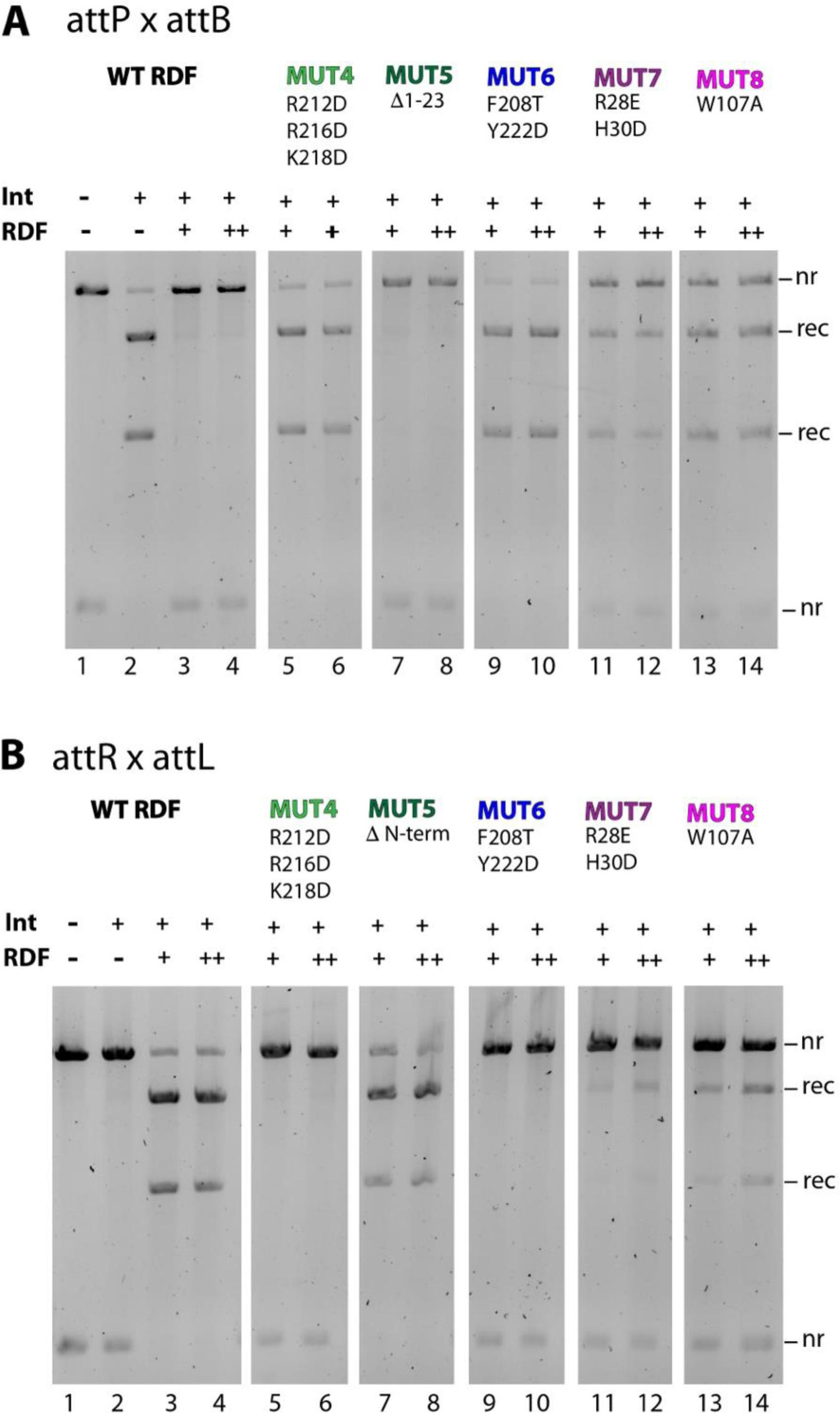
In vitro recombination activities of TG1 RDF mutants. Reactions were carried out for 2 hours in the reaction buffer described in the Methods section. **(A)** *attP* x *attB* recombination reactions. **(B)** *attR x attL* recombination reactions. Reaction products were digested with the restriction endonuclease NruI prior to 1.2% agarose gel electrophoresis. TG1 integrase (800 nM) was used in all reactions. Integrase was not added to reactions in lane 1 of A and B. The final RDF concentrations were 0 nM (-), 800nM (+) or 1600nM (++). The bands on the gel are labelled *nr* (non-recombinant, i.e., substrate) or *rec* (recombination product). The sizes of the products of the recombination reactions are shown alongside each band on the gel image. MUT4: TG1 RDF (R212D, R216D, K218D). MUT5: TG1 RDF (Δ1-23). MUT6: TG1 RDF (F208T, Y222D). MUT7: TG1 RDF (R28E H30D). MUT8: TG1 RDF (W107A).

First, we tested the activities of the mutant TG1 integrases in *attP* x *attB* reactions (which should be inhibited by the RDF) and in *attR* x *attL* reactions (which should be stimulated by the RDF) (Figure 6). Mutant 1 was less active than WT integrase: it showed limited activity on *attP* x *attB* after 2 hours (Figure 6A, lanes 2 vs. 4), but more noticeable activity after 16 hours (Figure 6B, lane 4). Given the overall defect of this mutant, it is unclear if there was an additional defect in RDF-mediated inhibition of its *attP* x *attB* reaction. However, despite showing some catalytic activity in the *attP* x *attB* reaction, mutant 1 completely failed to be stimulated by the RDF in an *attR* x *attL* reaction, even after 16 hours, indicating a defect in communication with the RDF (Figure 6C and D, lanes 5 and 6). Mutant 2, on the opposite side of DBD2 from mutant 1, showed near-WT activity in the *attP* x *attB* reaction (Figures 6A, lanes 2 vs. 7 and 6B lanes 2 vs. 7), but was defective in inhibition by the RDF (Figure 6B, lane 3 vs. 8 and 9). Furthermore, Mutant 2 was inefficiently stimulated by the RDF in the *attR* x *attL* reaction (Figure 6C and D, lane 3 vs. 8 and 9). Finally, the interactions of mutant 3 with the RDF could not be assessed because it was almost completely lacking in integrase activity (Figure 6B, lanes 10-12). This mutation was designed to test a more central region of the predicted interface, but the rather drastic change of two tryptophans to alanines probably destabilized the fold of DBD2.

Next we tested specific mutations in the TG1 RDF in similar assays. Mutations 4 and 6 showed the strongest effect – these RDFs completely failed to inhibit attP x attP recombination or to stimulate the *attR* x *attL* reaction (Figures 7A and B, lanes 3 and 4 vs. lanes 5, 6, 9 and 10). Figure 5E shows that they cluster with integrase mutation 2, at a point where the RDF is predicted to bind the DBD2-proximal segment of the coiled coil (pale green). Mutations 7 and 8 had a modest effect on RDF function (Figures 7A and B, lanes 3 and 4 vs. lanes 11-14). Figure 5D shows that these lie on the other side of the predicted interface. This result complements the findings that integrase mutation 1 at the same interface resulted in loss of RDF activity (Figures 5D and 6). Finally, RDF mutation 5 (Figure 5B, green), deletion of the N-terminal end (23 aa residues) had no effect on function (Figure 7A and B, lanes 3 and 4 vs. 7 and 8), an unsurprising result since that region of the RDF was predicted to be disordered and not in contact with the integrase (Figure 5C; only residue 23 is shown).

## DISCUSSION

In this study, we provide evidence that two different phage DNA replication proteins have been recruited to “moonlight” as RDFs for the serine integrase of the phage that encodes them. We experimentally verified that the TG1 RDF is a functional single-stranded DNA binding protein, and we found that the Bxb1 RDF is closely related structurally to an editing exonuclease. Our work strongly supports and expands previous proposals suggesting that recruitment of phage proteins for RDF function is a mechanism through which both LSIs and tyrosine-family integrases mediate excision reactions (2, 11, 12, 18).

Repurposing phage proteins to perform secondary functions allows frugal use of limited genome space, and is not limited to RDFs (44). For example, phage-iducible pathogenicity islands (PICIs) use phage proteins to moonlight as derepressors to lift repression of SapI induction. Examples of proteins used as PICI derepressors are dUTPase (SapIbov1 and SaPIbov5) (28); Sri, a phage protein involved in blocking bacterial DNA replication (SaPI1) (28); and DNA-single strand annealing proteins (SapI2) (45). Although moonlighting depressors and RDFs are similar in concept, the evolutionary paths to their repurposing may be different.

### Integrase proteins share an RDF-binding hotspot

The findings from this study, our previous modeling work (19), and the work of others all suggest that LSIs share a common RDF-binding hotspot, even though the RDFs do not share a common structure. Although there are no published experimental structures of integrase – RDF complexes, we can now use Alphafold to make testable predictions (19). In addition to the mutational testing of the TG1 – RDF interface model described here, we previously showed that alphafold2-multimer models are largely in agreement with prior mutational data for ϕC31, SPbeta, and Bxb1 (Supplementary Figure 2 of Shin *et al*., (19); (11, 41, 46). Previous work has also shown that the RDF redirects the coiled-coil, and that simply binding DBD2 is insufficient for function (42, 46, 47). We deduce that the RDF-binding hotspot lies at the DBD2-coiled coil junction, and includes the DBD2-proximal portion of the coiled coil. That portion was poorly ordered in the experimental structure of an LSI’s DNA binding domains with DNA (48) and is not predicted to be helical in all integrases, although we refer to it as part of the coiled coil for historical simplicity. We suggest that contacts to DBD2 itself may provide affinity for the RDF – integrase interaction, and that contacts to the DBD2-proximal coiled coil segment may be responsible for redirecting the trajectory of the coiled coil.

Here we tested a detailed Alphafold2-multimer model for the TG1 RDF – integrase interface. Although mutations across the predicted interface interfered with RDF function, the strongest effects were seen for mutations in the portion of the interface where the RDF is predicted to grip the DBD2-proximal portion of the integrase’s coiled coil (Figure 5E). Our finding that ssDNA interferes with TG1-RDF function *in vitro* (Figure 4) provides additional support for the model, because the positively charged surface on the RDF that is expected to bind ssDNA (Figure 2) is also used to bind the integrase (Figure 5A). These findings supprt the predicted the integrase-RDF interface, and suggest additional insights into how the RDF switches the directionality of the recombination reaction.

An intriguing feature not yet tested is the negatively charged, unstructured C-terminal tail of TG1 integrase – a feature not found in most LSIs, but shared by ϕC31 and ϕBT1 integrases, and others which use RDFs closely related to TG1’s (31). This tail may mimic ssDNA and compete with it for initial, transient binding to the RDF, which could lead to a high local concentration of RDF.

### How and why do phages moonlight existing proteins to take up RDF functions?

We hypothesize that the process of adopting a new RDF begins when a phage LSI becomes genetically separated from its old RDF (Figure 8). This could occur due to simultaneous activation of multiple prophages within a single host cell (49, 50). Recombination between replicating phage genomes could result in chimeric progeny encoding one parent phage’s LSI but not its cognate RDF. Such recombinant phage would be capable of infecting and lysogenizing a new host, but their further spread would be limited until they evolved to use another phage-encoded protein as an RDF. It is conceivable that most LSI RDFs could be scavenged proteins with or without known functions. It is likely that the original functions of some moonlighting RDFs have been taken over by other phage proteins while some retain their original function.

**Figure 8.**
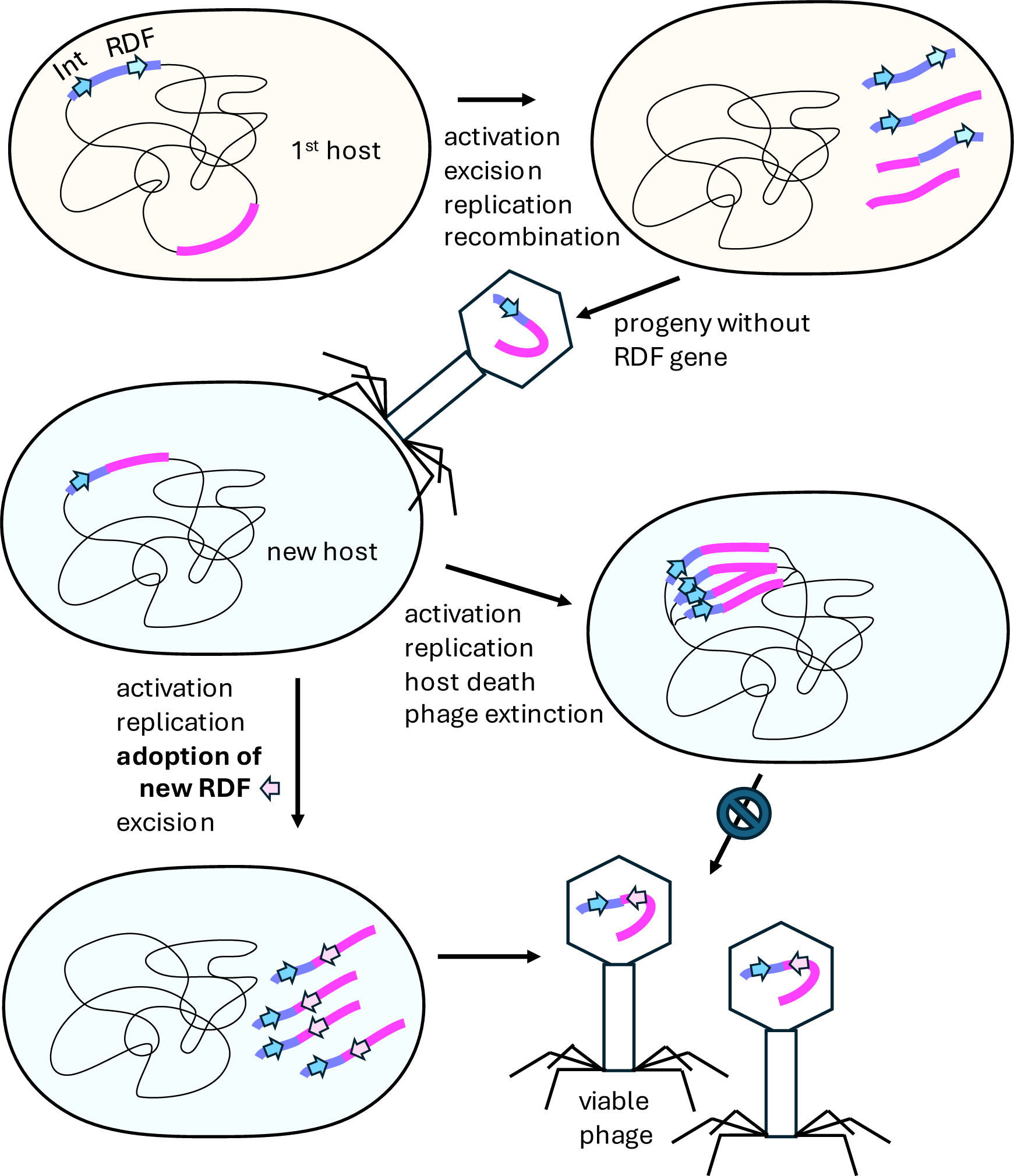
Proposed pathway for evolution of new RDFs. Recombination of two simultaneously activated prophages yields progeny with an integrase gene but no cognate RDF gene (top). The chimeric phage can establish a new lysogen (middle left) but is unable to excise from the host chromosome until it evolves to use another phage protein (pink arrow) as an RDF. The mutations that allow the use of a new RDF could occur during the prophage state or during replication.

The two previously identified RDFs studied in this work, those for Bxb1 and TG1, are both DNA replication proteins. Not only do they have the predicted structures of replication-related proteins, but the Bxb1 RDF is known to be required for phage replication as well as for excision (23) and in this work we showed that the TG1 RDF has ssDNA-binding activity. That some prophages can initiate replication before excision has been shown experimentally (51). However, a prophage that replicates within the host chromosome but never excises is likely to be an evolutionary dead end for both parties. Therefore, the integrase may be under strong selection for variants that can use one of the phage proteins already present as an RDF. This suggests that replication proteins may have been chosen because they were present in multiple copies at this make-or-break stage in the prophage life cycle. The existence of a binding hotspot on the integrase that is remote from its catalytic domain may facilitate such evolution.

The RDF binding hotspot may provide both advantages and disadvantages to the prophage. For example, an insufficiently selective integrase might bind constitutively expressed host proteins, triggering unregulated excision of the prophage. On the other hand, host proteins that are only expressed under conditions that induce prophage activation could be good candidates for moonlighting RDFs. In fact, such scavenging of host proteins could explain why we could not identify putative RDFs for all of the LSIs targets in our virtual pulldown study, which only considered phage-encoded proteins as possible RDFs (19). There is also precedent for crosstalk among mobile genetic elements: The RDF for some serine integrases found in PICI-like elements (PLEs) are not encoded by the PLE but come from the phages parasitised by the PLE (52).

The existence of a hotspot on LSIs where a second protein can bind, redirect the coiled-coil, and change the preferred reaction direction has implications for the evolution of RDFs and of directionality in LSIs in general. The original LSI may have been a simpler, bidirectional enzyme that simply used its coiled-coil subdomains to stabilize the synaptic complex that holds the two recombining DNAs together. For example, the LSIs encoded by the SCC*mec* family of mobile elements do not appear to follow the directionality “paradigm” of most characterized LSIs (53). Fortuitous interactions of the LSI hotspot with a second protein that could redirect the coiled coil’s trajectory may have initiated the evolutionary path toward control of directionality.

### Conclusion

In this report, we have shown that phage TG1 likely adapted its integrase to recognise a DNA single stranded binding protein as a cofactor for excisive recombination. Structural models of how TG1 integrase interacts with its RDF built using AlphaFold2 and AlphaFold multimer were validated through biochemical characterisation of integrase and RDF mutants. We also discuss strong evidence that the Bxb1 RDF is also a DNA replication protein, although a different one – an editing exonuclease. Future identification of the primary roles of other RDFs will rely on a high throughput version of AlphaFold Multimer that that can process virtual pulldowns on a metagenome-wide level. It will be interesting to know the mechanism through which phages select proteins to repurpose as RDFs.

## DATA AVAILABILITY

The data underlying this article are available in the article.

## AUTHOR CONTRIBUTIONS

Femi Olorunniji and Phoebe Rice: Conceptualization, Methodology, Writing, Funding acquisition. Abdulrazak Alsaleh, Tania Pena Reyes, Aron Baksh, Oluwateniola T. Taiwo-Aiyerin, Alexandria Holland, Ying Pigli: Methodology, Investigation.

## ACKNOWLEDGEMENTS

We thank Heewhan Shin and Adebayo Bello for helpful comments on the manuscript.

## REFERENCES

1. Jayaram, M., Ma, C.-H., Kachroo, A.H., Rowley, P.A., Guga, P., Fan, H.-F. and Voziyanov, Y. (2015) An Overview of Tyrosine Site-specific Recombination: From an Flp Perspective. Microbiol. Spectr., 3, 3.4.12.

2. Smith, M.C.M. (2015) Phage-encoded Serine Integrases and Other Large Serine Recombinases. Microbiol. Spectr., 3, 3.4.06.

3. Olorunniji, F.J., Rosser, S.J. and Stark, W.M. (2016) Site-specific recombinases: molecular machines for the Genetic Revolution. Biochem. J., 473, 673–684.

4. Grindley, N.D.F., Whiteson, K.L. and Rice, P.A. (2006) Mechanisms of site-specific recombination. Annu. Rev. Biochem., 75, 567–605.

5. Montano, S.P., Rowland, S.-J., Fuller, J.R., Burke, M.E., MacDonald, A.I., Boocock, M.R., Stark, W.M. and Rice, P.A. (2021) Structural basis for topological regulation of Tn3 resolvase. 10.1101/2021.12.07.471667.

6. Rowland, S., Boocock, M.R., Burke, M.E., Rice, P.A. and Stark, W.M. (2020) The protein– protein interactions required for assembly of the Tn *3* resolution synapse. Mol. Microbiol., 114, 952–965.

7. Mouw, K.W., Rowland, S.-J., Gajjar, M.M., Boocock, M.R., Stark, W.M. and Rice, P.A. (2008) Architecture of a serine recombinase-DNA regulatory complex. Mol. Cell, 30, 145– 155.

8. Laxmikanthan, G., Xu, C., Brilot, A.F., Warren, D., Steele, L., Seah, N., Tong, W., Grigorieff, N., Landy, A. and Van Duyne, G.D. (2016) Structure of a Holliday junction complex reveals mechanisms governing a highly regulated DNA transaction. eLife, 5, e14313.

9. Johnson, R.C. (2015) Site-specific DNA Inversion by Serine Recombinases. Microbiol. Spectr., 3, MDNA3-0047–2014.

10. Bibb, L.A., Hancox, M.I. and Hatfull, G.F. (2005) Integration and excision by the large serine recombinase phiRv1 integrase. Mol. Microbiol., 55, 1896–1910.

11. Ghosh, P., Wasil, L.R. and Hatfull, G.F. (2006) Control of phage Bxb1 excision by a novel recombination directionality factor. PLoS Biol., 4, e186.

12. Khaleel, T., Younger, E., McEwan, A.R., Varghese, A.S. and Smith, M.C.M. (2011) A phage protein that binds φC31 integrase to switch its directionality. Mol. Microbiol., 80, 1450–1463.

13. Van Duyne, G.D. and Rutherford, K. (2013) Large serine recombinase domain structure and attachment site binding. Crit. Rev. Biochem. Mol. Biol., 48, 476–491.

14. Van Duyne, G.D. (2001) A structural view of cre-loxp site-specific recombination. Annu. Rev. Biophys. Biomol. Struct., 30, 87–104.

15. Biswas, T., Aihara, H., Radman-Livaja, M., Filman, D., Landy, A. and Ellenberger, T. (2005) A structural basis for allosteric control of DNA recombination by λ integrase. Nature, 435, 1059–1066.

16. Landy, A. (2015) The λ Integrase Site-specific Recombination Pathway. Microbiol. Spectr., 3, MDNA3-0051–2014.

17. Van Duyne, G.D. and Landy, A. (2024) Bacteriophage lambda site-specific recombination. Mol. Microbiol., 121, 895–911.

18. Lewis, J.A. and Hatfull, G.F. (2001) Control of directionality in integrase-mediated recombination: examination of recombination directionality factors (RDFs) including Xis and Cox proteins. Nucleic Acids Res., 29, 2205–2216.

19. Shin, H., Holland, A., Alsaleh, A., Retiz, A.D., Pigli, Y.Z., Taiwo-Aiyerin, O.T., Reyes, T.P., Bello, A.J., Olorunniji, F.J. and Rice, P.A. (2024) Identification of cognate recombination directionality factors for large serine recombinases by virtual pulldown. 10.1101/2024.06.11.598349.

20. Jumper, J., Evans, R., Pritzel, A., Green, T., Figurnov, M., Ronneberger, O., Tunyasuvunakool, K., Bates, R., Žídek, A., Potapenko, A., et al. (2021) Highly accurate protein structure prediction with AlphaFold. Nature, 596, 583–589.

21. Evans, R., O’Neill, M., Pritzel, A., Antropova, N., Senior, A., Green, T., Žídek, A., Bates, R., Blackwell, S., Yim, J., et al. (2022) Protein complex prediction with AlphaFold-Multimer. 10.1101/2021.10.04.463034.

22. Abramson, J., Adler, J., Dunger, J., Evans, R., Green, T., Pritzel, A., Ronneberger, O., Willmore, L., Ballard, A.J., Bambrick, J., et al. (2024) Accurate structure prediction of biomolecular interactions with AlphaFold 3. Nature, 630, 493–500.

23. Savinov, A., Pan, J., Ghosh, P. and Hatfull, G.F. (2012) The Bxb1 gp47 recombination directionality factor is required not only for prophage excision, but also for phage DNA replication. Gene, 495, 42–48.

24. Yu, A. and Haggård-Ljungquist, E. (1993) The Cox protein is a modulator of directionality in bacteriophage P2 site-specific recombination. J. Bacteriol., 175, 7848–7855.

25. Esposito, D. and Scocca, J.J. (1994) Identification of an HP1 phage protein required for site-specific excision. Mol. Microbiol., 13, 685–695.

26. Saha, S., Haggård-Ljungquist, E. and Nordström, K. (1987) The cox protein of bacteriophage P2 inhibits the formation of the repressor protein and autoregulates the early operon. EMBO J., 6, 3191–3199.

27. Esposito, D., Wilson, J.C.E. and Scocca, J.J. (1997) Reciprocal Regulation of the Early Promoter Region of Bacteriophage HP1 by the Cox and CI Proteins. Virology, 234, 267–276.

28. Tormo-Más, M.Á., Mir, I., Shrestha, A., Tallent, S.M., Campoy, S., Lasa, Í., Barbé, J., Novick, R.P., Christie, G.E. and Penadés, J.R. (2010) Moonlighting bacteriophage proteins derepress staphylococcal pathogenicity islands. Nature, 465, 779–782.

29. Leveles, I., Németh, V., Szabó, J.E., Harmat, V., Nyíri, K., Bendes, Á.Á., Papp-Kádár, V., Zagyva, I., Róna, G., Ozohanics, O., et al. (2013) Structure and enzymatic mechanism of a moonlighting dUTPase. Acta Crystallogr. D Biol. Crystallogr., 69, 2298–2308.

30. Singh, M.I., Ganesh, B. and Jain, V. (2017) On the domains of T4 phage sliding clamp gp45: An intermolecular crosstalk governs structural stability and biological activity. Biochim. Biophys. Acta BBA - Gen. Subj., 1861, 3300–3310.

31. MacDonald, A.I., Baksh, A., Holland, A., Shin, H., Rice, P.A., Stark, W.M. and Olorunniji, F.J. (2024) Variable orthogonality of RDF - large serine integrase interactions within the ϕC31 family. BioRxiv Prepr. Serv. Biol., 10.1101/2024.04.03.587898.

32. Mirdita, M., Schütze, K., Moriwaki, Y., Heo, L., Ovchinnikov, S. and Steinegger, M. (2022) ColabFold - Making protein folding accessible to all. 10.1101/2021.08.15.456425.

33. Jurrus, E., Engel, D., Star, K., Monson, K., Brandi, J., Felberg, L.E., Brookes, D.H., Wilson, L., Chen, J., Liles, K., et al. (2018) Improvements to the APBS biomolecular solvation software suite. Protein Sci., 27, 112–128.

34. Holm, L., Laiho, A., Törönen, P. and Salgado, M. (2023) DALI shines a light on remote homologs: One hundred discoveries. Protein Sci., 32, e4519.

35. Sauguet, L., Raia, P., Henneke, G. and Delarue, M. (2016) Shared active site architecture between archaeal PolD and multi-subunit RNA polymerases revealed by X-ray crystallography. Nat. Commun., 7, 12227.

36. Bianco, P.R. (2022) OB-fold Families of Genome Guardians: A Universal Theme Constructed From the Small β-barrel Building Block. Front. Mol. Biosci., 9, 784451.

37. Shamoo, Y., Friedman, A.M., Parsons, M.R., Konigsberg, W.H. and Steitz, T.A. (1995) Crystal structure of a replication fork single-stranded DNA binding protein (T4 gp32) complexed to DNA. Nature, 376, 362–366.

38. Abioye, J., Lawson-Williams, M., Lecanda, A., Calhoon, B., McQue, A.L., Colloms, S.D., Stark, W.M. and Olorunniji, F.J. (2023) High fidelity one-pot DNA assembly using orthogonal serine integrases. Biotechnol. J., 18, e2200411.

39. Olorunniji, F.J., McPherson, A.L., Rosser, S.J., Smith, M.C.M., Colloms, S.D. and Stark, W.M. (2017) Control of serine integrase recombination directionality by fusion with the directionality factor. Nucleic Acids Res., 45, 8635–8645.

40. Olorunniji, F.J., Lawson-Williams, M., McPherson, A.L., Paget, J.E., Stark, W.M. and Rosser, S.J. (2019) Control of ϕC31 integrase-mediated site-specific recombination by protein trans-splicing. Nucleic Acids Res., 47, 11452–11460.

41. Abe, K., Takahashi, T. and Sato, T. (2021) Extreme C-terminal element of SprA serine integrase is a potential component of the ‘molecular toggle switch’ which controls the recombination and its directionality. Mol. Microbiol., 115, 1110–1121.

42. Mandali, S., Gupta, K., Dawson, A.R., Van Duyne, G.D. and Johnson, R.C. (2017) Control of Recombination Directionality by the Listeria Phage A118 Protein Gp44 and the Coiled-Coil Motif of Its Serine Integrase. J. Bacteriol., 199, e00019–17.

43. Bibb, L.A. and Hatfull, G.F. (2002) Integration and excision of the Mycobacterium tuberculosis prophage-like element, phiRv1. Mol. Microbiol., 45, 1515–1526.

44. Penadés, J.R. and Christie, G.E. (2015) The Phage-Inducible Chromosomal Islands: A Family of Highly Evolved Molecular Parasites. Annu. Rev. Virol., 2, 181–201.

45. Bowring, J., Neamah, M.M., Donderis, J., Mir-Sanchis, I., Alite, C., Ciges-Tomas, J.R., Maiques, E., Medmedov, I., Marina, A. and Penadés, J.R. (2017) Pirating conserved phage mechanisms promotes promiscuous staphylococcal pathogenicity island transfer. eLife, 6, e26487.

46. Fogg, P.C.M., Younger, E., Fernando, B.D., Khaleel, T., Stark, W.M. and Smith, M.C.M. (2018) Recombination directionality factor gp3 binds ϕC31 integrase via the zinc domain, potentially affecting the trajectory of the coiled-coil motif. Nucleic Acids Res., 46, 1308–1320.

47. Rowley, P.A., Smith, M.C.A., Younger, E. and Smith, M.C.M. (2008) A motif in the C-terminal domain of ϕC31 integrase controls the directionality of recombination. Nucleic Acids Res., 36, 3879–3891.

48. Rutherford, K., Yuan, P., Perry, K., Sharp, R. and Van Duyne, G.D. (2013) Attachment site recognition and regulation of directionality by the serine integrases. Nucleic Acids Res., 41, 8341–8356.

49. De Paepe, M., Hutinet, G., Son, O., Amarir-Bouhram, J., Schbath, S. and Petit, M.-A. (2014) Temperate Phages Acquire DNA from Defective Prophages by Relaxed Homologous Recombination: The Role of Rad52-Like Recombinases. PLoS Genet., 10, e1004181.

50. Dragoš, A., Priyadarshini, B., Hasan, Z., Strube, M.L., Kempen, P.J., Maróti, G., Kaspar, C., Bose, B., Burton, B.M., Bischofs, I.B., et al. (2021) Pervasive prophage recombination occurs during evolution of spore-forming *Bacilli*. ISME J., 15, 1344–1358.

51. Chen, J., Quiles-Puchalt, N., Chiang, Y.N., Bacigalupe, R., Fillol-Salom, A., Chee, M.S.J., Fitzgerald, J.R. and Penadés, J.R. (2018) Genome hypermobility by lateral transduction. Science, 362, 207–212.

52. McKitterick, A.C. and Seed, K.D. (2018) Anti-phage islands force their target phage to directly mediate island excision and spread. Nat. Commun., 9, 2348.

53. Misiura, A., Pigli, Y.Z., Boyle-Vavra, S., Daum, R.S., Boocock, M.R. and Rice, P.A. (2013) Roles of two large serine recombinases in mobilizing the methicillin-resistance cassette SCCmec. Mol. Microbiol., 88, 1218–1229.

